# Adult porcine intestinal organoids as models for regional epithelial identity and individual regulatory variation

**DOI:** 10.1101/2025.10.29.679705

**Authors:** Fany Blanc, Smahane Chalabi, Frederic Pepke, Mayrone Mongellaz, Mathieu Charles, Andrea Rau, Sarah Djebali, Giorgia Egidy, Elisabetta Giuffra

## Abstract

**Background:** Organoids offer controlled epithelial systems for functional follow-up of candidate genes and variants in farm animals, but their usefulness for genotype-to-phenotype research depends on whether they retain intestinal regional identity and animal-specific molecular signatures. We profiled 32 RNA-seq libraries from matched tissues and organoids derived from duodenum, jejunum, ileum and colon of four slaughter-age Large White pigs.

**Results:** Organoids were transcriptionally distinct from native tissues but remained within the intestinal transcriptional space and retained regional structure, particularly when epithelial cell signature genes were considered. Most expressed genes were shared between tissues and organoids (81.2% of all detected genes and 81.8% of epithelial cell signature genes). Innate immune genes were also largely detected in organoids (79.6% shared), but with a lower regional organisation. Projection of ileal organoids onto an *in vivo* developmental reference positioned them closer to foetal or newborn samples than to slaughter-age ileum tissue, consistent with a comparatively immature transcriptional state. Differential expression and enrichment analyses showed attenuation of gene-level segment differences in organoids but partial preservation of regional markers and Gene Ontology terms linked to intestinal development and regionalisation, absorption, metabolism, epithelial signalling, mucosal immune responses and transport. Animal-associated differences were retained for selected epithelial signatures, notably the glycosylation-related genes *FUT2* and *B4GALNT2*. *FUT2* expression was strongly reduced in one animal and associated with linked intronic variants, whereas *B4GALNT2* separated two pairs of animals and segregated with candidate variants affecting protein expression or transcript processing.

**Conclusions:** Adult porcine intestinal organoids retain a substantial epithelial component of regional identity and preserve selected animal-specific regulatory signatures, while showing reduced immune spatial patterning and immature transcriptional features. Standardised organoid biobanks from genotyped and phenotyped pigs may support functional follow-up of candidate genes, variants and pathways when intestinal epithelial mechanisms are implicated.

## Background

Understanding how genetic variation shapes cellular pathways underlying complex traits remains a central challenge in farmed animal genetics. Advanced predictive models coupled to deep genome annotations and multi-omics approaches are increasingly able to nominate candidate variants, regulatory mechanisms and cellular pathways at scale, but causal validation remains a major bottleneck. In this context, advanced *in vitro* models, including organoids, are increasingly viewed as a useful experimental interface between genomic prediction and animal phenotypes, with additional benefits for the 3Rs (Replacement, Reduction and Refinement) principles in animal research [1–3].

In farmed species, organoid-based platforms are increasingly being developed for tissue-specific functional studies such as host–microbe interactions, nutritional or infectious challenges [4,5]. Adult stem cell-derived intestinal organoids reproduce key properties of the native intestinal epithelium, including epithelial organisation, barrier function, absorption and metabolic activity [6,7]. In this study, we focused on two features that are relevant for the use of these models in genotype-to-phenotype research: the retention of segment-specific epithelial identity and the preservation of animal-specific molecular signatures that may reflect genetic or regulatory variation.

Several studies have benchmarked porcine intestinal organoids against their tissue of origin. Jejunal organoids have been shown to retain transcriptomic similarity to native epithelium and to remain stable during long-term culture [8]. Organoids derived from jejunum and colon of suckling and weaned piglets preserved gut-region signatures more consistently than developmental-stage signatures [9]. More recently, ileal and colonic organoids from pigs divergent for feed efficiency were used to investigate transcriptomic differences and responses to *Escherichia coli*, supporting the potential of porcine intestinal organoids for studies related to production and health traits [10]. These studies established porcine intestinal organoids as robust epithelial models, but they left open the extent to which organoids derived from multiple adult gut segments retain both regional transcriptional programmes and animal-specific regulatory signatures within the same experimental design. In humans, donor-to-donor variability in intestinal organoids has been shown to remain detectable across functional readouts [11], and donor-specific *ACE2* expression in intestinal organoid-derived monolayers has been linked to differences in susceptibility to SARS-CoV-2 infection [12]. Whether comparable animal-specific transcriptional signatures are preserved in porcine intestinal organoids, and whether they can highlight candidate regulatory variation, remains insufficiently explored. This question is relevant because several complex traits in farm animals, including nutrient use, feed efficiency, host defence and resilience, involve components linked to intestinal epithelial function. Standardised organoid biobanks derived from genotyped and phenotyped animals could provide controlled cellular models for functional follow-up of candidate genes, variants and pathways [2,3].

We previously established standardised procedures for generating, cryopreserving and expanding organoids from duodenum, jejunum, ileum and colon of Large White pigs at slaughter age, and showed that these organoids retain segment-associated morphological and cellular features [13]. Here, we characterise the transcriptome of intestinal tissues and matched organoids from the same four animals and four gut segments. We assessed global and epithelial transcriptional identity, compared innate immune signatures, projected ileal organoids onto an *in vivo* developmental reference, and analysed regional and between-animal expression differences. Finally, we explored candidate variants at two glycosylation-related loci, *FUT2* and *B4GALNT2*, as examples of animal-specific regulatory signatures retained *in vitro*.

## Methods

### Tissues, organoid cultures, RNA extractions

Intestinal samples were obtained from four Large White pigs at slaughter age from the INRAE Experimental Unit UE3P (Saint Gilles, France) in the context of routine commercial slaughter. Duodenum, jejunum, ileum and colon samples were collected from each animal. For each gut segment, a native tissue sample was preserved for RNA extraction and a matched sample was used to establish 3D intestinal organoids. Protocols and metadata for tissue sampling, organoid generation, culture and RNA isolation have been deposited on the Functional Annotation of Animal Genomes portal [14]. In brief, 3D organoids were generated, cryopreserved at passage 2 and expanded in IntestiCult™ Organoid Growth Medium (Human) (STEMCELL Technologies), according to previously described procedures [4,13], with minor modifications for cryopreservation in FBS/DMSO [15] (Supplementary Figure 1A). When using frozen biobanked organoids to establish batches, the most consistent procedures were achieved after three cycles of subculture in the early passages. RNA was isolated from intestine segments (duodenum, jejunum, ileum, and colon) tissues and their derived organoids [15]. To minimise technical variation, library preparation and sequencing were performed in a single experimental batch. High-quality RNA was obtained for all tissue samples (mean RIN 7.5, range 6.5–8.0) and organoid samples (mean RIN 9.7, range 8.3–10.0).

### RNA-seq libraries and sequencing

Libraries were prepared from 200 ng of total RNA and sequenced as 2 x 101-bp paired-end reads on an Illumina NovaSeq 6000 system using a NovaSeq 6000 S4 Reagent Kit (200 cycles) at the iGenSeq transcriptomics platform (Brain and Spine Institute, Paris, France). To account for the cellular complexity and heterogeneity of intestinal tissues, strand-specific poly(A)+ RNA-seq libraries were sequenced to a high depth, with approximately 2 × 60 million paired-end reads per biological replicate, to support detection of low-abundance and tissue-specific transcripts. Quality control confirmed that all 32 RNA-seq libraries were suitable for downstream analyses (Supplementary Table 1). Power analysis using RNASeqPower indicated >80% power to detect two-fold expression differences with four biological replicates at FDR < 0.05, given typical RNA-seq variance estimates.

Raw data and associated metadata are available through the FAANG Data Portal [14].

### RNA-seq data processing and gene annotation

RNA-seq data were processed using the TAGADA pipeline v2.0.0, which performs read mapping, transcript reconstruction and expression quantification [16]. Reads were aligned to the pig reference genome Sscrofa11.1, Ensembl release 108, using STAR [17], and transcript expression was quantified with StringTie [18]. Downstream analyses were performed in R [19]. Gene names were annotated using BioMart v2.54.0 and the HCOP orthology database [20] and are provided in Supplementary Table 2. Normalised TPM values were filtered to retain genes with TPM > 0.1 in at least two samples and log2-transformed for downstream visualisation and correlation analyses.

Multidimensional scaling (MDS) was used to visualise transcriptome-wide relationships among tissues and organoids across gut segments. To compare gene detection between tissues and organoids, a gene was defined as expressed when its TPM value exceeded the selected threshold in at least 75% of replicates within a given condition. Genes were then classified as shared when detected in both tissues and organoids, tissue-specific when detected only in tissues, or organoid-specific when detected only in organoids. This analysis was performed both across all samples and within each intestinal segment. The overall transcriptomic analysis workflow is shown in Supplementary Figure 1B.

### Visualisation and comparison with public data

To assess whether organoids retained an intestinal transcriptional identity in a broader tissue context, RNA-seq profiles from the present study were analysed together with publicly available GENE-SWitCH transcriptomic datasets from liver, muscle and duodenum collected from the same four animals (BioProject PRJEB58031; [21]). These GENE-SWitCH datasets were processed using the same TAGADA-based workflow described above before joint visualisation. This comparison was used for multidimensional scaling analysis and is shown in Supplementary Figure 2.

To evaluate the relationship between ileal organoids and *in vivo* intestinal maturation, ileum tissues and ileum-derived organoids from the present study were analysed together with the GENE-SWitCH ileum tissue datasets representing foetal day 30 (F30), foetal day 70 (F70) and newborn piglets (NB) [22]. PCA was performed using all expressed genes, epithelial cell signature genes and innate immune genes. For each gene set, centroids for F30, F70 and NB samples were used to define an *in vivo* developmental trajectory, and the centroids of slaughter-age ileum tissues and ileum-derived organoids were projected orthogonally onto this trajectory.

Epithelial cell signature genes were defined using porcine ileum single-cell RNA-seq data from Wiarda et al. [23], selecting genes from clusters 36 and 45, corresponding to epithelial cells. Innate immune genes were defined using InnateDB v5.4 [24].

### Differential expression and enrichment analyses

Differential expression analyses were conducted independently for tissues and organoids samples using limma-voom framework and the edgeR Bioconductor package (v3.40.0). Read counts were normalised using the calcNormFactors function and Limma’s voom function (v3.54.0) was then used to fit a single linear model including gut segment and animal as fixed effects. The proportions of variance explained by each of these factors were evaluated using the fitExtractVarPartModel function of the variancePartition package (v1.28.9). Differential expression associated with either gut segment or animal was then assessed from this models while accounting for the other factor. P-values were adjusted using the false discovery rate (FDR) method, and FDR < 0.05 was considered significant. Feature set enrichment analyses were performed using the tmod R package (v0.50.13). The Coincident Extreme Ranks in Numerical Observations (CERNO) method was used to identify enriched terms from ranked lists of annotated human orthologs based on logFC values from the differential expression analyses, using decreasing ranks for upregulated genes and increasing ranks for downregulated genes. Gene Ontology Biological Process terms were obtained from msigdbr v7.5.1 and used for functional interpretation. Enrichment was considered significant at FDR < 0.05.

### Variant identification and annotation

Whole-genome sequence data for the four animals were obtained from the GENE-SWitCH project [25]. Animals SSC_INRAE_GS_WP4_97W to SSC_INRAE_GS_WP4_100W are here referred to as A1 to A4. Variants located in the genomic regions encompassing the annotated *FUT2* and B4GALNT2 gene bodies and extending 2 kb upstream of their transcription start sites were extracted and annotated using the Variant Effect Predictor v111.1 [26]. Splice-affecting variants, including variants located in intronic or splice-region contexts, were predicted using Pangolin v1.0.1 [27,28]. The GENCODE-like GTF annotation file for Sus scrofa used as input for Pangolin was generated from Ensembl GTF files. This approach has previously been applied to non-human mammals [29].

## Results

### Organoids are transcriptionally distinct from tissues but retain intestinal and epithelial regional identity

Organoids were generated, cryopreserved and expanded according to previously established standards for this sample collection (11; see Materials and Methods). We generated 32 RNA-seq libraries from matched intestinal tissues and organoids spanning duodenum, jejunum, ileum, and colon in four Large White pigs. Transcriptome quantification identified a total of 19,929 expressed genes, with an average of 18,666 genes detected per sample.

As a first broader comparison, intestinal organoid and tissue transcriptomes were analysed together with liver, muscle and duodenum transcriptomes from the same four animals. Data visualisation by multidimensional scaling (MDS) confirmed that, as expected, organoids were clearly separated from native tissues along the first dimension, which explained 33% of total variance. However, organoid transcriptomes clustered with intestinal samples rather than with non-intestinal tissues from the same animals, supporting the maintenance of an intestinal transcriptional identity *in vitro* (Supplementary Figure 2).

When only organoid and tissue samples from the four gut segments were analysed (Figure 1A), the first MDS dimension separated organoids from tissues explaining 55% of total variance while 9% of total variance in the second dimension reflected intestinal regional origin. Native tissues clustered distinctly by gut segment, whereas organoids showed a clear separation between small-intestinal and colonic samples, with partial discrimination among the three small-intestinal regions. These results indicate that, despite major transcriptomic differences between tissues and organoids, intestinal organoids retain in part the regional component of their tissue of origin.

**Figure 1.**
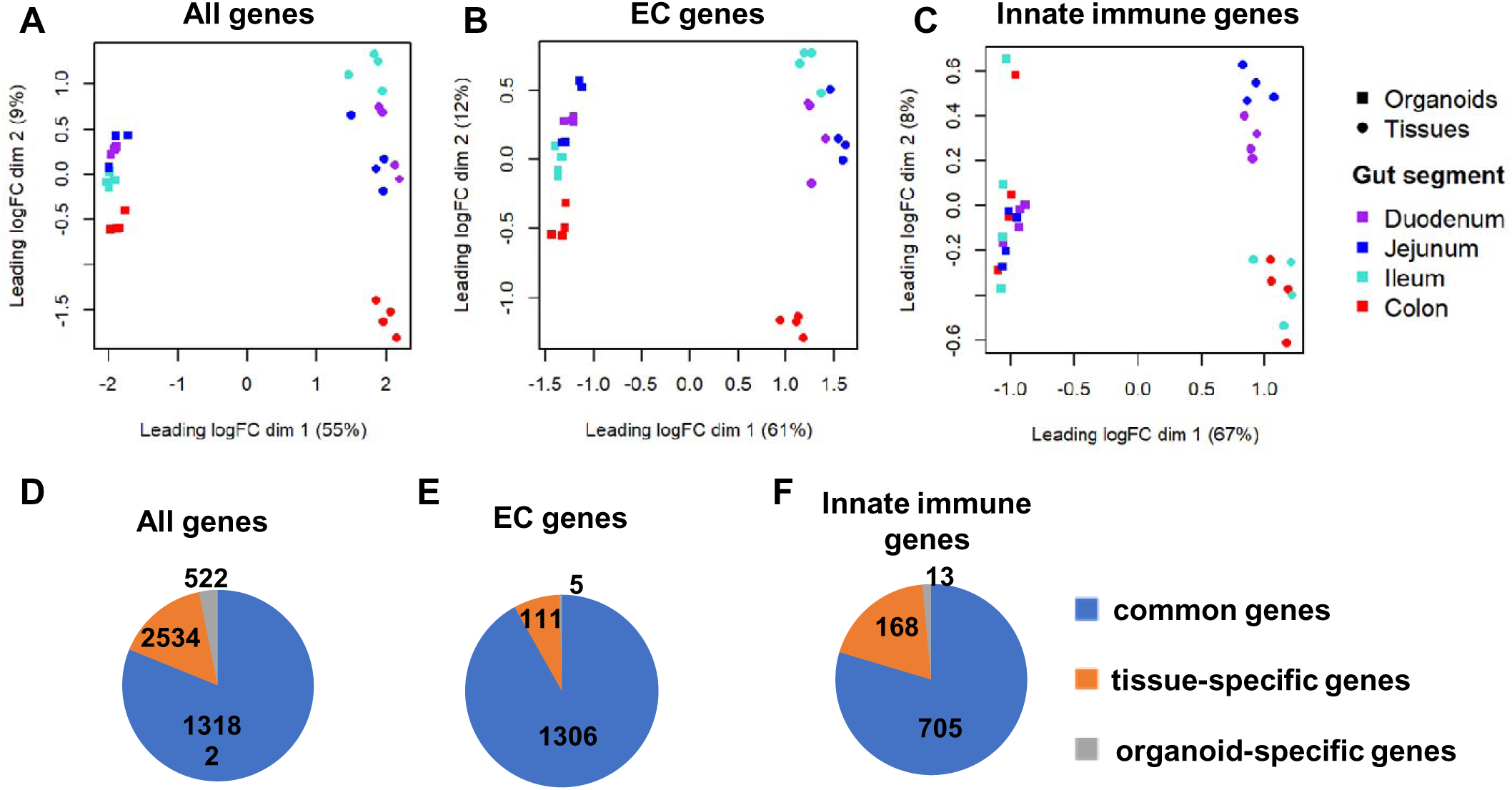
Transcriptomic comparison of matched intestinal tissues and organoids. A–C. Multidimensional scaling analysis of intestinal tissues and matched organoids using: A, all expressed genes; B, epithelial cell signature genes; and C, innate immune genes. D–F. Number of genes detected in both tissues and organoids, detected only in tissues, or detected only in organoids for: D, all expressed genes; E, epithelial cell signature genes; and F, innate immune genes. Genes were considered expressed when TPM exceeded the defined threshold in the required number of replicates. Epithelial cell signature genes were derived from porcine ileum single-cell RNA-seq data, and innate immune genes were defined using InnateDB. EC, epithelial cell; MDS, multidimensional scaling; TPM, transcripts per million.

To determine how this regional structure was retained within the epithelial compartment, we focused on an epithelial cell signature derived from porcine ileum single-cell RNA-seq data [23]. MDS analysis based on 1,431 expressed epithelial cell signature genes reproduced the separation between tissues and organoids along the first dimension and retained regional organisation along the second dimension (Figure 1B). This pattern indicates that epithelial transcriptional programmes remain regionally patterned in organoids.

We then compared genes detected in intestinal tissues and their matched organoids. Across all gut segments, most expressed genes were shared between tissues and organoids, corresponding to 81.2% of the detected transcriptome, whereas 15.6% were detected only in tissues and 3.2% only in organoids (Figure 1D). Similar proportions were observed when each intestinal segment was analysed separately, with shared genes ranging from 79.6% to 82.5% (Supplementary Figure 3A).

Among epithelial cell signature genes, the overlap between tissues and organoids was high, with 81.8% shared genes, 7.8% tissue-specific genes and 0.4% organoid-specific genes (Figure 1E). This pattern remained stable across gut segments and across TPM thresholds ranging from 0.1 to 10 (Supplementary Figure 3A,B). To assess expression-level concordance beyond gene detection, we computed Spearman correlations of log2-transformed TPM values between paired tissue and organoid samples across gut regions and animals. Correlations ranged from 0.53 to 0.88, indicating that matched organoids retain part of the transcriptional hierarchy observed in their tissues of origin while also showing the expected divergence associated with an *in vitro* epithelial culture (Supplementary Figure 3D).

### Innate immune signatures are reduced and less spatially resolved in organoids

Because native intestinal tissues contain immune, stromal and other non-epithelial cell populations that are not represented in epithelial organoid cultures, we next examined genes associated with innate immune responses separately. MDS analysis based on 958 expressed innate immune genes (InnateDB resource) showed that duodenum and jejunum tissues segregated from ileum and colon tissues, whereas organoids clustered more closely together (Figure 1C). This indicates that regional patterning of innate immune-related transcription is less pronounced in organoids than in native tissues.

Consistent with this observation, most innate immune genes detected in tissues were also detected in organoids, but the fraction of tissue-specific genes was higher than for epithelial cell signature genes. Overall, 79.6% of innate immune genes were shared between tissues and organoids, 19.0% were detected only in tissues and 1.4% only in organoids (Figure 1F). Thus, although organoids retain a substantial fraction of innate immune-related transcripts, their spatial resolution across intestinal regions is reduced compared with native tissues.

### Developmental trajectory analysis suggests an immature transcriptional state in ileal organoids

To infer how ileal organoids relate to *in vivo* intestinal maturation, we analysed ileum tissue and organoid transcriptomes vs. the GENE-SWitCH ileum datasets spanning two foetal stages, at 30- and 70-days post-fertilisation (F30 and F70), and newborn piglets (NB). This analysis was used to project organoid transcriptomes onto an *in vivo* developmental reference rather than to assign them to a precise developmental stage.

PCA based on all expressed genes revealed a transcriptional trajectory associated with intestinal maturation (Figure 2A). F30, F70 and NB samples aligned progressively along the first principal component (PC1), consistent with a developmental continuum. Ileum tissue samples collected at slaughter age extended this trajectory towards a more mature transcriptional state. In contrast, ileum organoids were displaced from the main *in vivo* trajectory, although their orthogonal projections onto the developmental axis were positioned between F70 and NB samples.

**Figure 2.**
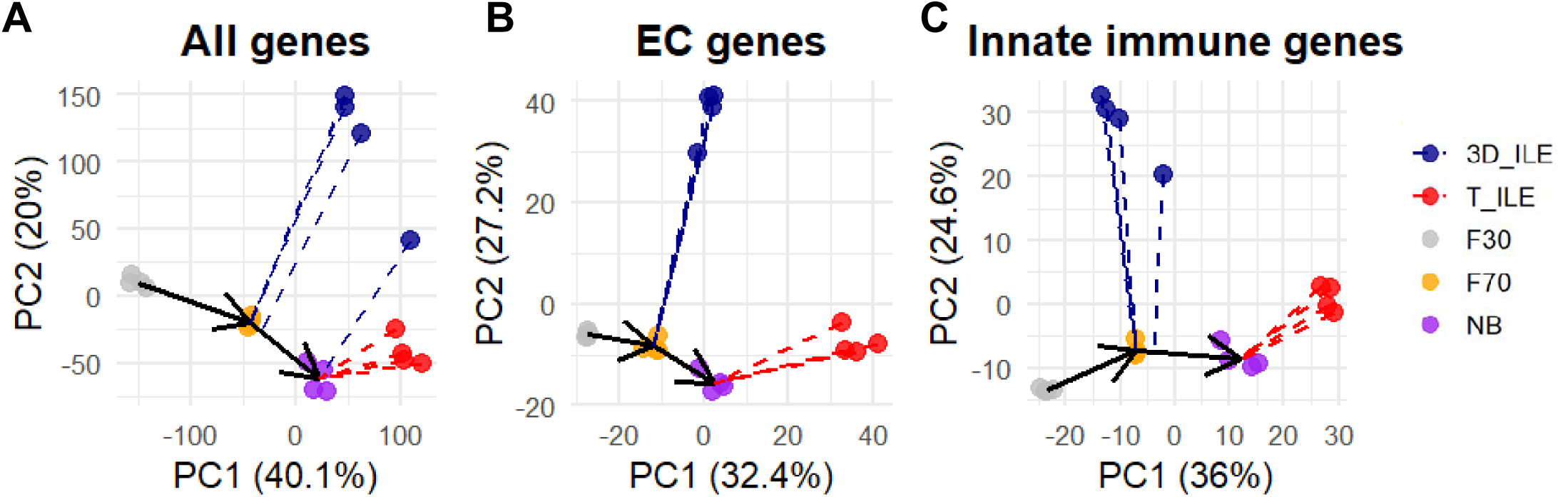
Developmental projection of ileal organoids onto an in vivo *ileum reference.* PCA of ileum tissues and ileum-derived organoids together with GENE-SWitCH ileum tissue datasets representing foetal day 30 (F30), foetal day 70 (F70) and newborn piglets (NB). Analyses were performed using A, all expressed genes; B, epithelial cell signature genes; and C, innate immune genes. The F30, F70 and NB samples define an *in vivo* developmental trajectory, which is extended by slaughter-age ileum tissue samples. Ileal organoids are displaced from this main trajectory, but their orthogonal projections fall closer to foetal or newborn developmental stages than to adult ileum tissue, consistent with a comparatively immature transcriptional state. PC, principal component; F30, foetal day 30; F70, foetal day 70; NB, newborn; T_ILE, ileum tissue; 3D_ILE, ileum organoids.

When PCA was restricted to epithelial cell and innate immune gene sets (Figure 2B,C), ileum tissue samples collected at slaughter age again positioned along the developmental trajectory. Ileum organoids remained outside the principal developmental axis, but their orthogonal projections were closer to F70 samples. This pattern indicates that the immature-like transcriptional state of organoids is detectable not only at the whole-transcriptome level but also within epithelial and innate immune-related gene subsets.

Together, these analyses suggest that porcine ileal organoids retain transcriptional features associated with intermediate or immature developmental programmes rather than fully recapitulating adult ileum tissue.

### Regional functional programmes are partially preserved despite reduced gene-level differentiation

To quantify the main sources of transcriptomic variation, we first partitioned gene expression variance according to gut segment and animal. In tissue samples, an average of 51.7% of the observed variance was attributed to gut segment, whereas 17.1% was attributed to animal. In organoids, the proportion of variance explained by gut segment was lower, at 31.0%, whereas the contribution of animal effects increased to 27.4%. Thus, regional origin remained a major determinant of gene expression, but animal-associated variation became comparatively more prominent in organoids.

Differential expression analyses between gut segments confirmed that regional transcriptional differences were attenuated in organoids compared with native tissues. Across pairwise segment comparisons, tissues showed between 3,279 and 8,711 differentially expressed genes (DEGs; FDR < 0.05), whereas organoids showed between 39 and 1,858 DEGs for the same comparisons (Table 1; Supplementary Table 2). Although the number of DEGs was markedly lower in organoids, a subset of genes was consistently differentially expressed in both tissues and organoids for the same segment comparisons. The number of shared DEGs ranged from 7 for ILE vs. JEJ to 596 for COL vs. DUO (Table 1).

**Table 1.**
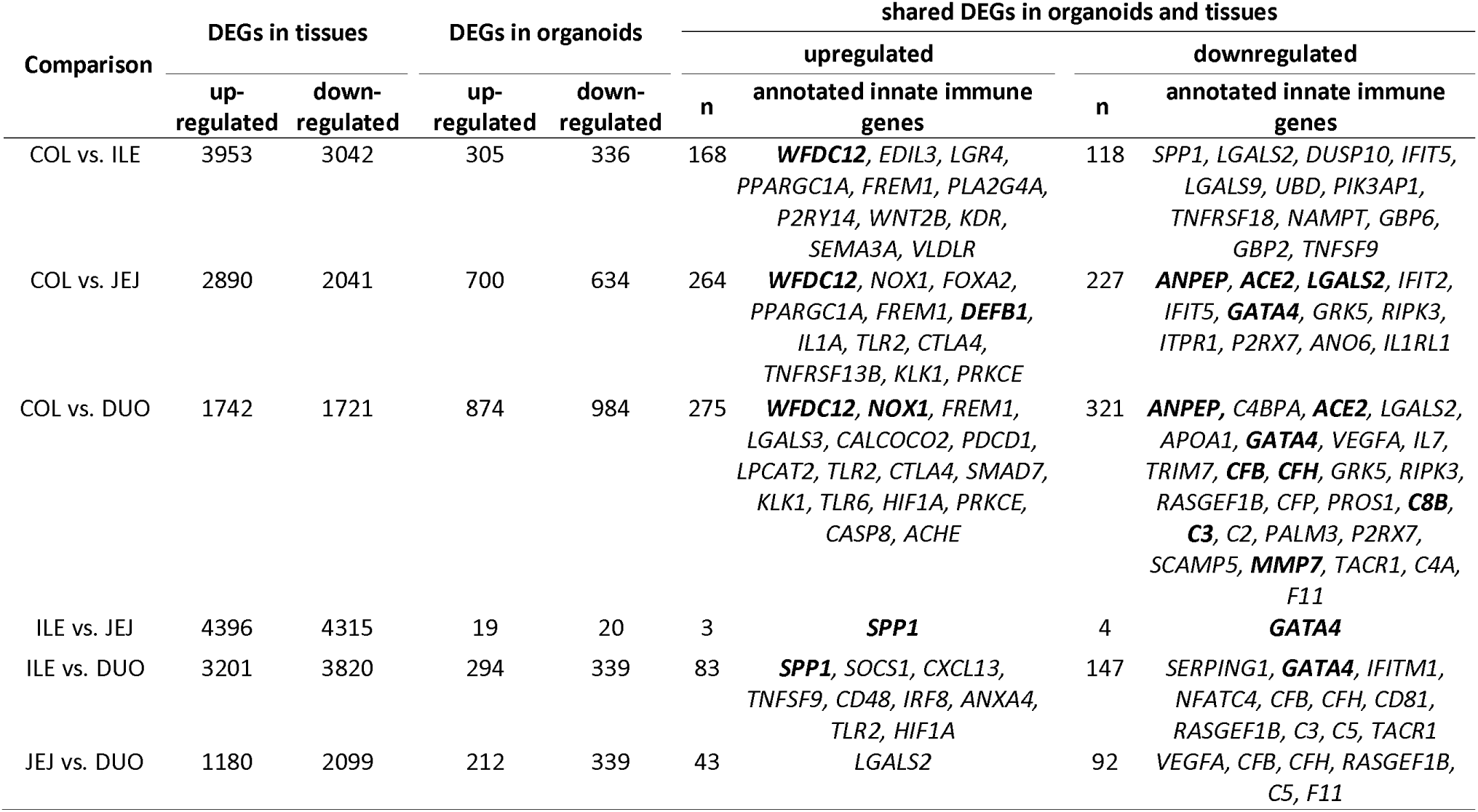
Differentially expressed genes across intestinal segments in tissues and organoids.

Notably, these shared DEGs included functionally informative regional markers. In comparisons between colon and small-intestinal segments, colon samples showed higher expression of genes such as *WFDC12, FREM1, NOX1* and *TLR2*, whereas small-intestinal samples were enriched for markers including *LGALS2, GATA4* and *ACE2*. Within the small intestine, shared regional differences included higher expression of complement-related genes such as *C3* and *CFB*, together with *VEGFA*, in duodenum; higher *SPP1* and *CXCL13* and lower *GATA4* in ileum; and higher *LGALS2* in jejunum relative to adjacent segments. These shared expression patterns support partial conservation of biologically meaningful segment-associated transcriptional programmes in organoids, despite the overall reduction in DEG numbers.

For each pairwise comparison, the table reports the number upregulated and downregulated differentially expressed genes (DEGs) identified between intestinal segments in tissues and organoids. Under “shared DEGs in organoids and tissues”, the table reports the total number of DEGs identified in both tissue and organoid analyses with the same direction of regulation and the annotated innate immune genes among these shared DEGs. Genes shown in bold exhibited an absolute log₂ fold change > 1 in both tissue and organoid analyses. DEGs were defined using FDR < 0.05. DUO, duodenum; JEJ, jejunum; ILE, ileum; COL, colon.

To gain functional insight into these regional differences, we performed feature set enrichment analysis (FSEA) using log₂FC values from the differential expression analyses. In tissues, FSEA identified between 339 and 1,378 significantly enriched GO terms across segment comparisons, whereas in organoids 400 to 995 enriched GO terms were detected (Supplementary Table 3; Supplementary Figures 4 and 5). We then compared organoid and tissue FSEA results to identify biological processes shared between corresponding segment contrasts.

Shared enriched GO terms were detected across all segment comparisons, ranging from 84 terms for JEJ vs. DUO to 278 for COL vs. ILE (Figure 3; Supplementary Table 3). These shared terms were associated with major functional programmes contributing to intestinal compartmentalisation, including intestinal development and regionalisation, digestive and absorptive functions, xenobiotic and metabolic adaptation, mucosal immune responses, epithelial signalling and ion or molecular transport homeostasis. Many of these terms were enriched in the same direction in organoids and tissues, supporting preservation of regional functional hallmarks despite reduced gene-level differentiation *in vitro*. Some tissue-associated metabolic and absorptive processes, particularly in comparisons involving ileum and jejunum, were less consistently detected in organoids.

**Figure 3.**
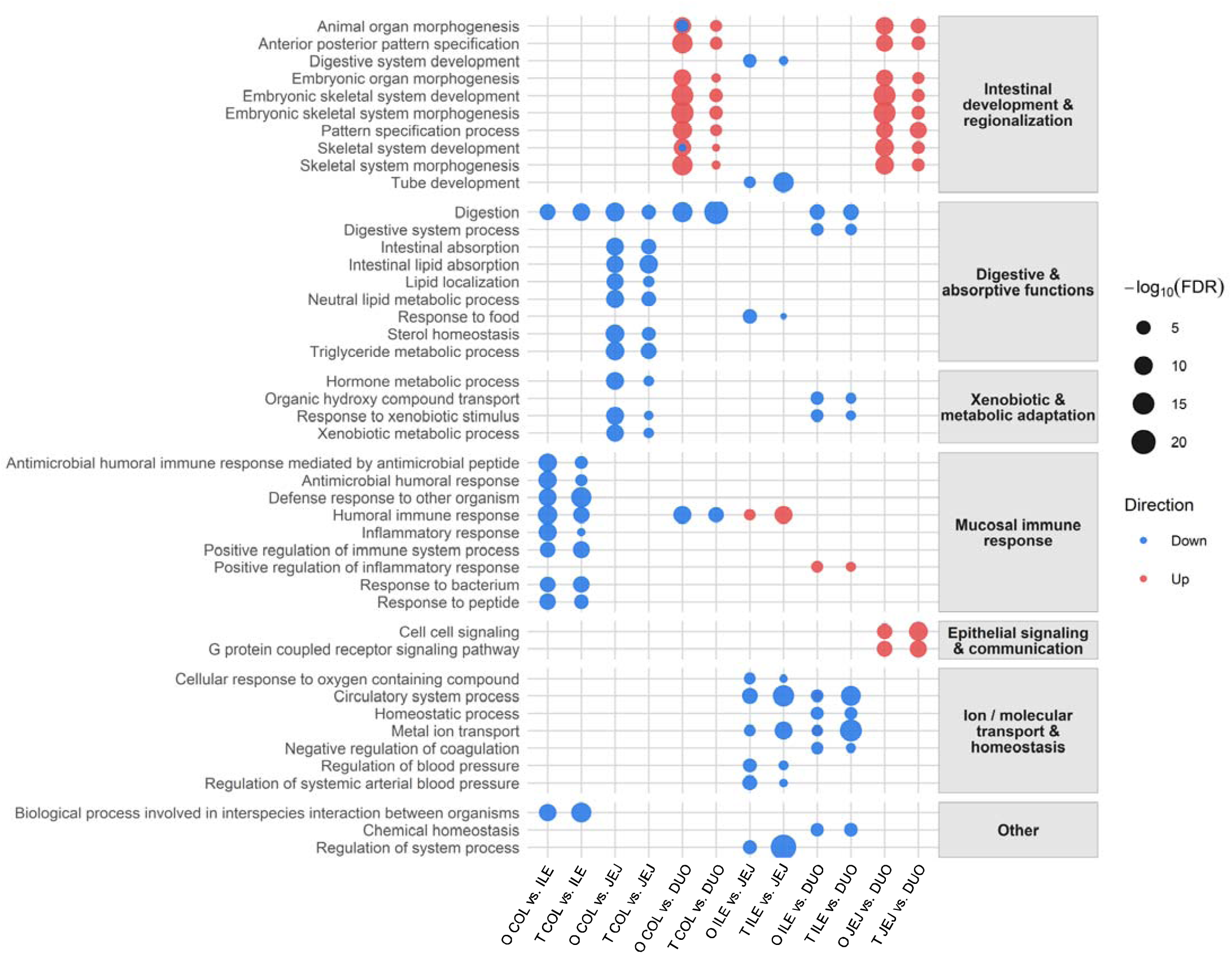
Regional functional programmes shared between tissues and organoids. Functional categorisation of Gene Ontology (GO) terms enriched in organoid segment comparisons and also detected in the corresponding tissue comparisons. Feature set enrichment analysis (FSEA) was performed using ranked log₂FC values from pairwise differential expression analyses between intestinal segments. For each segment comparison, the figure shows enriched GO terms shared between organoids and matched tissues, grouped into major biological categories. Dot size represents –log₁₀(FDR), and dot colour indicates the direction of regulation in the corresponding comparison. O, organoid; T, tissue; DUO, duodenum; JEJ, jejunum; ILE, ileum; COL, colon.

### Organoids retain selected animal-specific glycosylation-related signatures

Differential expression analyses were also performed between individual animals, in parallel for tissues and organoids. The number of animal-associated DEGs varied markedly across pairwise comparisons and between the two systems. In tissues, 52 to 731 DEGs were identified across animal comparisons, whereas in organoids 4 to 1,396 DEGs were detected (FDR < 0.05; Table 2; Supplementary Table 4). Table 2 summarizes the numbers of up- and downregulated DEGs between animals in tissues and organoids and highlights their overlap.

**Table 2.**
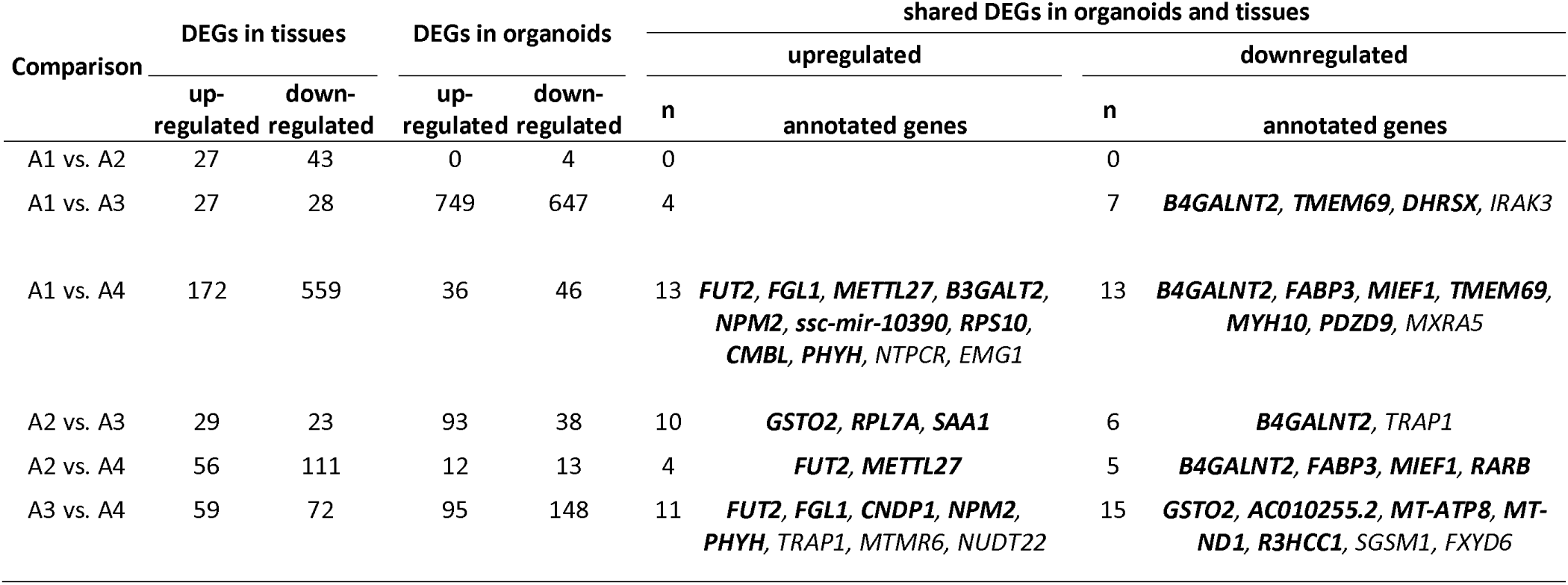
Differentially expressed genes (DEGs) identified in pairwise comparisons between animals in tissues and organoids.

Only a limited number of genes were consistently up- or downregulated in both tissues and organoids for the same animal comparisons, ranging from 0 for A1 vs. A2 to 26 for A1 vs. A4 and A3 vs. A4 (Table 2). Among these shared animal-associated differences, the clearest signals involved genes related to epithelial glycosylation. *B4GALNT2* consistently differentiated A1 and A2 from A3 and A4 in both tissues and organoids (Figure 4A). *FUT2* showed a marked A4-specific reduction in expression (Figure 4B), together with other genes including *FGL1*, *METTL27*, *NPM2* and *PHYH* (Supplementary Figure 6A–D). Additional animal-associated expression patterns included higher *FABP3* expression in A3 and A4, higher *TRAP1* and *GSTO2* expression in A3, and lower *TMEM69* expression in A1 (Supplementary Figure 6E–H). These results indicate that organoids retain selected animal-specific transcriptional signatures observed in native tissues, with the strongest evidence provided by *FUT2* and *B4GALNT2*.

**Figure 4.**
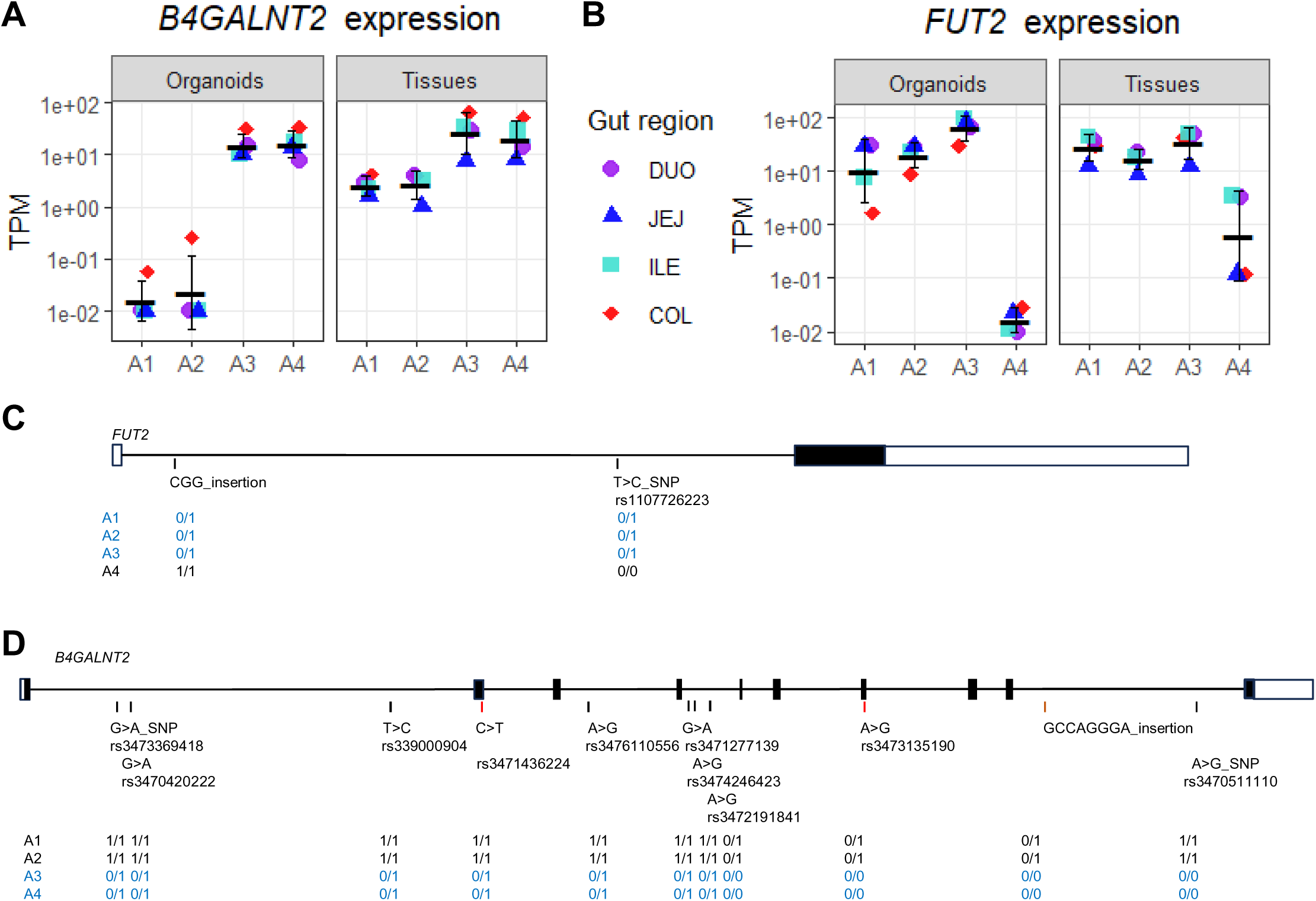
Animal-specific expression of *B4GALNT2* and *FUT2* variants segregating with expression patterns. A–B. Expression levels of *B4GALNT2* and *FUT2* in intestinal tissues and matched organoids from four animals across duodenum, jejunum, ileum and colon. Expression is shown as TPM. C–D. Gene structures and variants identified in the FUT2 and B4GALNT2 genomic regions using whole-genome sequence data from the same animals. Genotypes are shown for each animal after comparison with the observed expression patterns. *FUT2* expression was markedly reduced in animal A4, particularly in organoids. *B4GALNT2* expression separated animals A1/A2 from A3/A4 in both tissues and organoids. These patterns are consistent with candidate cis-regulatory variation affecting glycosylation-related gene expression. DUO, duodenum; JEJ, jejunum; ILE, ileum; COL, colon; TPM, transcripts per million.

For each pairwise comparison, the table reports the number of upregulated and downregulated DEGs identified between animals in tissues and organoids. The numbers under “shared DEGs in organoids and tissues” indicate the total number of DEGs identified in both tissue and organoid analyses with the same direction of regulation. Annotated genes among these shared DEGs are listed. DEGs were defined using FDR < 0.05. Genes shown in bold exhibited an absolute log₂ fold change > 1 in both tissue and organoid analyses. A1–A4, individual animals.

The marked inter-individual differences observed for *FUT2* and *B4GALNT2* expression suggested the presence of underlying candidate cis-regulatory variation (Figure 4A,B). To investigate this hypothesis, we analysed whole-genome sequence data from the four animals [25] and searched for variants segregating with the observed expression patterns.

For *FUT2*, animals A1–A3 showed substantially higher expression levels than A4 in both tissues and organoids, although the difference was more pronounced *in vitro*. In intestinal tissues, *FUT2* expression ranged from 18 to 39 TPM in A1–A3 but dropped to 1.7 TPM in A4. In organoids, expression was nearly abolished in A4 (0.016 TPM), whereas A1–A3 ranged from 17 to 70 TPM, representing a >200-fold difference (Figure 4A). Genomic analysis identified two linked intronic variants defining two *FUT2* haplotypes: a reference REF-T haplotype and an alternative CGG-C haplotype carrying a CGG trinucleotide insertion in intron 1 together with a linked SNP (Figure 4C; Supplementary Table 5). Animal A4 was homozygous for the alternative haplotype, whereas A1–A3 were heterozygous. Although these variants were not predicted to alter splicing, their segregation with *FUT2* expression is consistent with a candidate regulatory effect.

For *B4GALNT2*, animals A1 and A2 consistently displayed low expression levels, whereas A3 and A4 showed markedly higher expression in both tissues and organoids (Figure 4B). Variant screening across the *B4GALNT2* locus identified candidate segregating variants associated with this expression pattern, including variants predicted to modify protein expression or transcript processing (Figure 4D; Supplementary Tables 6 and 7). Transcript-level analysis indicated that the animal-specific *B4GALNT2* expression pattern reflected a locus-level difference rather than a switch between major isoforms (Supplementary Figure 7; Supplementary Table 8).

Together, these results indicate that intestinal organoids can retain and, in some cases, amplify selected animal-specific epithelial expression signatures observed in native tissues. *FUT2* and *B4GALNT2* provide candidate examples of glycosylation-related regulatory variation that is preserved *in vitro*.

## Discussion

Adult porcine intestinal organoids provide controlled epithelial models that can help bridge functional genomics and genotype-to-phenotype studies in pigs. Using matched tissues and organoids from four gut segments and four slaughter-age pigs, we show that adult-derived organoids retain a substantial epithelial component of intestinal regional identity and preserve selected animal-specific epithelial expression signatures, while also revealing model boundaries related to innate immune spatial patterning and maturation state.

Previous studies have established porcine intestinal organoids as epithelial models, showing transcriptomic similarity to native epithelium, partial retention of gut-region signatures and applicability to health- and production-related questions [8–10]. Our study extends these observations by assessing organoids derived from four adult intestinal segments within a matched tissue–organoid design, and by distinguishing regional epithelial programmes from innate immune-related signatures and animal-associated expression differences. Although organoids were globally distinct from native tissues, they remained positioned within the intestinal transcriptomic space, particularly when epithelial cell signature genes were considered (Figure 1A,B,D,E; Supplementary Figures 2 and 3). This is consistent with single-cell studies showing epithelial regional specialisation along the porcine intestine [23,30]. Thus, adult-derived porcine intestinal organoids do not represent generic intestinal epithelial cultures, but preserve a biologically meaningful subset of segment-associated epithelial programmes.

This epithelial regionalisation was reflected in differential expression and pathway enrichment analyses. Although the number of segment-associated DEGs was lower in organoids than in native tissues, shared DEGs were detected across all segment comparisons and included functionally informative markers of intestinal compartmentalisation (Table 1; Supplementary Table 2). Colon–small intestine contrasts retained differences involving genes such as *NOX1*, *TLR2*, *GATA4*, *LGALS2* and *ACE2*, whereas small-intestinal comparisons highlighted regional markers including *C3*, *CFB*, *VEGFA*, *SPP1*, *CXCL13* and *LGALS2*. These patterns are consistent with known regional differences in epithelial defence, absorptive function, signalling and mucosal immune communication along the intestine [30–33]. The partial retention of these regional markers in organoids therefore supports the preservation of an epithelial component of intestinal functional zonation. At the pathway level, shared enriched GO terms involved intestinal development and regionalisation, digestive and absorptive functions, xenobiotic and metabolic adaptation, epithelial signalling, mucosal immune-related responses and ion or molecular transport homeostasis (Figure 3; Supplementary Table 3). Thus, standard porcine intestinal organoids preserve a coherent subset of segment-associated epithelial programmes, making them particularly relevant for testing hypotheses in which the primary expected effect of genetic or epigenetic variation is epithelial-centred.

Among animal-associated expression differences, *FUT2* and *B4GALNT2* provided the clearest examples of signatures shared between native tissues and organoids (Figure 4A,B). These signatures were not only retained *in vitro*, but in some cases appeared more pronounced in organoids, consistent with enrichment of epithelial cell-centred programmes in this system. This is biologically relevant because *FUT2* and *B4GALNT2* encode functionally related glycosyltransferases acting on partly overlapping glycan substrates in intestinal epithelial cells. Their coordinated or reciprocal activity contributes to mucosal glycosylation patterns, with major implications for host–microbe interactions, susceptibility to infectious disease and immune homeostasis [34,35]. At *FUT2*, linked intronic variants segregated with the strong reduction of expression observed in animal A4, without predicted effects on splicing (Figure 4C). At *B4GALNT2*, segregating variants were associated with contrasting expression profiles between animals, including variants predicted to affect protein expression or transcript processing, while transcript-level analyses did not support a major isoform switch (Figure 4D). The fact that these expression phenotypes were preserved, and sometimes amplified, in organoids lacking microbiota and non-epithelial cellular complexity suggests that at least part of the underlying regulation is epithelial-intrinsic, potentially reflecting cis-regulatory effects, epigenetic features retained from the tissue of origin [36], or a combination of both.

These observations do not identify causal variants, but provide a proof-of-principle that matched porcine organoid systems can retain, and in some cases reveal more clearly, animal-specific epithelial expression differences. This is particularly relevant when genetic or phenotypic evidence points to the intestinal epithelium as a candidate tissue involved in an *in vivo* trait. In such cases, organoids can provide a controlled model to prioritise candidate regulatory mechanisms and to design subsequent functional validation, including perturbation, genome editing or allele-specific assays [3,37,38].

These findings also clarify two relevant model boundaries. First, although most innate immune genes detected in tissues were also expressed in organoids, their regional organisation was attenuated *in vitro* (Figure 1C,F). Immune-related transcription in native intestinal tissue is not uniform along the gut and does not follow a single cephalocaudal gradient; instead, each intestinal segment has a distinct immune gene-expression profile, as shown in human and mouse studies [33]. The reduced spatial resolution observed in organoids therefore suggests that basal immune-related transcriptional patterns are only partly retained and are disconnected from the multicellular tissue environment that contributes to segment-specific immune identity *in vivo*. Second, after projection onto the GENE-SWitCH *in vivo* developmental reference, ileal organoids projected closer to foetal or newborn reference samples, both at the whole-transcriptome level and when epithelial or innate immune-related genes were analysed separately (Figure 2A–C). While this analysis cannot assign a precise developmental stage to these organoids, it confirms that these organoids retain an immature transcriptional component. This feature is consistent with observations that intestinal organoids can preserve developmental or maturation-related cues, and should be considered when modelling pathways influenced by epithelial differentiation, immune maturation or interactions with the native tissue environment [39].

Future genotype-to-phenotype applications will therefore depend on matching model complexity to the biological question and the required experimental scale. Standard intestinal epithelial organoids are best positioned as scalable systems for the functional follow-up of candidate genes, variants and pathways emerging from farm-animal G2P studies, particularly when the expected effect is epithelial-centred [2,3]. By contrast, mechanistic dissection of immune–epithelial crosstalk, stromal, vascular or neuronal regulation, or host–microbiota interactions will require complexified organoids, organ-on-chip platforms or other advanced *in vitro* configurations [3–5].

## Conclusions

Adult porcine intestinal organoids derived from multiple gut segments retain a substantial component of intestinal and epithelial regional identity despite being transcriptionally distinct from native tissues and comparatively immature. They also preserve selected animal-specific epithelial expression signatures, including glycosylation-related differences involving *FUT2* and *B4GALNT2*, which may be associated with candidate regulatory variation. At the same time, standard epithelial organoids show reduced regional organisation of innate immune-related signatures and retain an immature transcriptional component, highlighting the need to match model complexity to the biological question. These findings support the use of standardised porcine intestinal organoid biobanks from genotyped and phenotyped animals as controlled systems for functional follow-up of candidate genes, variants and pathways, particularly when *in vivo* evidence points to the intestinal epithelium as a relevant tissue for the trait.

## Declarations

### Ethics approval and consent to participate

The Large White pigs analysed in this study were reared and slaughtered at the INRAE UE3P phenotyping facility [40] in the context of the EU-H2020 GENE-SWitCH project. Intestinal samples were obtained post mortem from animals slaughtered in the context of routine production and phenotyping activities. No live-animal experimental procedures were performed specifically for the purpose of this study. Therefore, specific ethical approval for animal experimentation was not required for the present work, in accordance with the European Union Directive 2010/63/EU on the protection of animals used for scientific purposes. Consent to participate was not applicable.

### Consent for publication

Not applicable.

### Availability of data and materials

The RNA-seq data and associated metadata generated for this study are available through the FAANG Data Portal under BioProject PRJEB64165 [14], under the FAANG Data Sharing Statement. Public GENE-SWitCH transcriptomic datasets used for comparison with liver, muscle and duodenum samples are available through the FAANG Data Portal under BioProject PRJEB58031 [21]. Public GENE-SWitCH ileum transcriptomic datasets used for developmental projection analyses are available under BioProject PRJEB41970 [22].

### Funding

This study was supported by GENE-SWitCH, a project funded by the European Union’s Horizon 2020 Research and Innovation Programme under grant agreement No. 817998, and by INRAE, Department of Animal Genetics, through the SNOOPY and IMMUNOID projects.

Whole-genome sequence data used for variant analyses are available under BioProject PRJEB58030 [25]. Supplementary tables and figures are provided as additional files.

### Authors’ contributions

FB, GE and EG conceived and designed the study. EG secured funding. FB, FP and MM prepared samples and generated data. FB, SC, AR, SD, MC, GE and EG analysed and interpreted the data. FB, SC, GE and EG drafted the manuscript. FB, GE and EG revised and finalised the manuscript. All authors read and approved the final manuscript.

## Acknowledgements

The authors thank the INRAE UE3P phenotyping facility for animal management and sampling support, and the INRAE @Bridge platform for use of the equipment for RNA extraction and quality control. This work benefited from equipment and services from the iGenSeq core facility at the Brain and Spine Institute, ICM, Paris, France.

---

Additional file 1_Supplementary **Tables 1–8**. File format: .xlsx.

**Supplementary Table 1:** Characteristics of the 32 RNA-seq datasets from pig tissues and organoids and quality control.

**Supplementary Table 2:** Differential analysis results comparing the different gut segments in tissues and organoids.

**Supplementary Table 3:** Feature Set Enrichment Analysis to evaluate GO:BP terms enriched in differentially expressed genes comparing the different gut segments in tissues and organoids.

**Supplementary Table 4:** Differential analysis results comparing the different animals in tissues and organoids.

**Supplementary Table 5:** Predicted functional impact of variants segregating with *FUT2* expression levels on splicing in the four animals.

**Supplementary Table 6:** Predicted functional impact of variants segregating with *B4GALNT2* expression levels on splicing in the four animals.

**Supplementary Table 7:** Predicted effect of variants with modifying power on *B4GALNT2* expression.

**Supplementary Table 8**: Expression of individual *B4GALNT2* isoforms in the four animals.

**Additional file 2_Supplementary Figures 1–7**. File format: .pdf.

**Supplementary Figure 1:** Experimental design and transcriptomic analysis workflow. Overview of sample collection, organoid generation, RNA-seq data production and downstream transcriptomic analyses. Intestinal tissues and matched organoids were obtained from four Large White pigs at slaughter age across four gut segments. The workflow includes global transcriptomic comparisons, segment-wise differential expression, functional enrichment analysis, animal-specific expression analyses and candidate variant investigation using whole-genome sequence data.

**Supplementary Figure 2:** Global positioning of intestinal organoids relative to intestinal and non-intestinal tissues. Multidimensional scaling analysis including intestinal organoids, matched intestinal tissues and additional tissue transcriptomes from the same animals. Organoids separate from native tissues along the first dimension but cluster with intestinal samples rather than with non-intestinal tissues, supporting the maintenance of an intestinal transcriptional identity.

**Supplementary Figure 3:** Robustness of gene-sharing patterns between tissues and organoids. A. Number of genes detected in both tissues and organoids, only in tissues, or only in organoids, for each intestinal segment. B. Sensitivity analysis showing how the number of shared and sample-type-specific genes changes across TPM thresholds. C. Distribution of expression levels in tissues and organoids across intestinal segments. D. Spearman correlations between matched tissue and organoid samples across animals and intestinal regions.

**Supplementary Figure 4:** Tissue-based regional functional enrichment. Gene Ontology Biological Process terms enriched in tissue segment comparisons based on feature set enrichment analysis of ranked log₂FC values. The top enriched upregulated and downregulated GO:BP terms are shown for each tissue segment comparison. Dot size represents –log₁₀(FDR), and dot colour indicates direction of regulation.

**Supplementary Figure 5:** Organoid-based regional functional enrichment. Gene Ontology Biological Process terms enriched in organoid segment comparisons based on feature set enrichment analysis of ranked log₂FC values. The top enriched upregulated and downregulated GO:BP terms are shown for each organoid segment comparison. Dot size represents –log₁₀(FDR), and dot colour indicates direction of regulation.

**Supplementary Figure 6:** Animal-associated expression patterns for selected genes. Expression levels of selected genes showing animal-associated differences in tissues and organoids. For each animal, expression across the four intestinal segments is shown separately for tissues and organoids.

**Supplementary Figure 7:** *B4GALNT2* isoform expression across animals and gut segments. Expression levels of *B4GALNT2* transcript isoforms across intestinal tissues and matched organoids from the four animals. The dominant isoform accounts for most *B4GALNT2* expression in both tissues and organoids, indicating that the animal-associated expression pattern reflects a locus-level difference rather than a major isoform switch.

## References

1. Clark EL, Archibald AL, Daetwyler HD, Groenen MAM, Harrison PW, Houston RD, et al. From FAANG to fork: application of highly annotated genomes to improve farmed animal production. Genome Biol. 2020;21:285. 10.1186/s13059-020-02197-8

2. Clark EL, Amaral A, Baranasic D, Barta E, Cartick G, Chua P, et al. Establishing the ELIXIR Domestic Animals Genome and Phenome Community. F1000Res. 2026;15:339. 10.12688/f1000research.177500.1

3. Giuffra E, Alberio R, Bickhart D, Boyartchuk V, Brevini TAL, Carpi D, et al. In Vitro Systems for Genotype-to-Phenotype Research in Farmed Animals: From Functional Genomic Screens to Mechanistic Insights [Internet]. Biology and Life Sciences; 2026 [cited 2026 July 29]. 10.20944/preprints202607.1543.v1

4. Beaumont M, Blanc F, Cherbuy C, Egidy G, Giuffra E, Lacroix-Lamandé S, et al. Intestinal organoids in farm animals. VETERINARY RESEARCH. 2021;52. 10.1186/s13567-021-00909-x

5. Kwon D-H, Kwon H, Jang G. Organoid-based platforms in livestock: Current advances and future prospects. Research in Veterinary Science. 2026;198:105985. 10.1016/j.rvsc.2025.105985

6. Sato T, Vries RG, Snippert HJ, van de Wetering M, Barker N, Stange DE, et al. Single Lgr5 stem cells build crypt-villus structures in vitro without a mesenchymal niche. Nature. 2009;459:262–5. 10.1038/nature07935

7. Rossi G, Manfrin A, Lutolf MP. Progress and potential in organoid research. Nat Rev Genet. 2018;19:671–87. 10.1038/s41576-018-0051-9

8. Van Der Hee B, Madsen O, Vervoort J, Smidt H, Wells JM. Congruence of Transcription Programs in Adult Stem Cell-Derived Jejunum Organoids and Original Tissue During Long-Term Culture. Front Cell Dev Biol. 2020;8:375. 10.3389/fcell.2020.00375

9. Mussard E, Lencina C, Gallo L, Barilly C, Poli M, Feve K, et al. The phenotype of the gut region is more stably retained than developmental stage in piglet intestinal organoids. Front Cell Dev Biol. 2022;10:983031. 10.3389/fcell.2022.983031

10. Madsen O, Rikkers RSC, Wells JM, Bergsma R, Kar SK, Taverne N, et al. Transcriptomic analysis of intestinal organoids, derived from pigs divergent in feed efficiency, and their response to Escherichia coli. BMC Genomics. 2024;25:173. 10.1186/s12864-024-10064-0

11. Mohammadi S, Morell-Perez C, Wright CW, Wyche TP, White CH, Sana TR, et al. Assessing donor-to-donor variability in human intestinal organoid cultures. Stem Cell Reports. 2021;16:2364–78. 10.1016/j.stemcr.2021.07.016

12. Jang KK, Kaczmarek ME, Dallari S, Chen Y-H, Tada T, Axelrad J, et al. Variable susceptibility of intestinal organoid–derived monolayers to SARS-CoV-2 infection. Koo B-K, editor. PLoS Biol. 2022;20:e3001592. 10.1371/journal.pbio.3001592

13. Blanc F, Mongellaz M, Pepke F, Bevilacqua C, Riviere J, Vilotte M, et al. 505. Phenotypic characterization of organoids derived from pig intestine segments. Proceedings of 12th World Congress on Genetics Applied to Livestock Production (WCGALP) [Internet]. Leiden, The Netherlands: Brill; 2025 [cited 2026 July 29]. p. 2097–100. 10.3920/978-90-8686-940-4_505

14. PRJEB64165 | FAANG dataset [Internet]. [cited 2026 July 30]. https://data.faang.org/dataset/PRJEB64165. Accessed 30 July 2026

15. INRAE_SOP_generating_organoids_20230329.pdf [Internet]. [cited 2026 July 31]. https://api.faang.org/files/protocols/samples/INRAE_SOP_generating_organoids_20230329.pdf. Accessed 31 July 2026

16. Kurylo C, Guyomar C, Foissac S, Djebali S. TAGADA: a scalable pipeline to improve genome annotations with RNA-seq data. NAR Genomics and Bioinformatics. 2023;5:lqad089. 10.1093/nargab/lqad089

17. Dobin A, Davis CA, Schlesinger F, Drenkow J, Zaleski C, Jha S, et al. STAR: ultrafast universal RNA-seq aligner. Bioinformatics. 2013;29:15–21. 10.1093/bioinformatics/bts635

18. Pertea M, Kim D, Pertea GM, Leek JT, Salzberg SL. Transcript-level expression analysis of RNA-seq experiments with HISAT, StringTie and Ballgown. Nat Protoc. 2016;11:1650–67. 10.1038/nprot.2016.095

19. Ihaka R, Gentleman R. R: A Language for Data Analysis and Graphics. Journal of Computational and Graphical Statistics. 1996;5:299–314. 10.1080/10618600.1996.10474713

20. HCOP: Orthology Predictions Search | HUGO Gene Nomenclature Committee [Internet]. [cited 2026 July 30]. https://www.genenames.org/tools/hcop/. Accessed 30 July 2026

21. PRJEB58031 | FAANG dataset [Internet]. [cited 2026 July 31]. https://data.faang.org/dataset/PRJEB58031. Accessed 31 July 2026

22. PRJEB41970 | FAANG dataset [Internet]. [cited 2026 July 30]. https://data.faang.org/dataset/PRJEB41970. Accessed 30 July 2026

23. Wiarda JE, Trachsel JM, Sivasankaran SK, Tuggle CK, Loving CL. Intestinal single-cell atlas reveals novel lymphocytes in pigs with similarities to human cells. Life Sci Alliance. 2022;5:e202201442. 10.26508/lsa.202201442

24. InnateDB: Systems Biology of the Innate Immune Response [Internet]. [cited 2026 July 30]. https://www.innatedb.com/annotatedGenes.do?type=innatedb. Accessed 30 July 2026

25. PRJEB58030 | FAANG dataset [Internet]. [cited 2026 July 30]. https://data.faang.org/dataset/PRJEB58030. Accessed 30 July 2026

26. McLaren W, Gil L, Hunt SE, Riat HS, Ritchie GRS, Thormann A, et al. The Ensembl Variant Effect Predictor. Genome Biol. 2016;17:122. 10.1186/s13059-016-0974-4

27. Zeng T, Li YI. Predicting RNA splicing from DNA sequence using Pangolin. Genome Biol. 2022;23:103. 10.1186/s13059-022-02664-4

28. Zeng T. tkzeng/Pangolin [Internet]. 2026 [cited 2026 July 31]. https://github.com/tkzeng/Pangolin. Accessed 31 July 2026

29. Charles M, Gaiani N, Sanchez M-P, Boussaha M, Hozé C, Boichard D, et al. Functional impact of splicing variants in the elaboration of complex traits in cattle. Nat Commun. 2025;16:3893. 10.1038/s41467-025-58970-5

30. Wiarda JE, Becker SR, Sivasankaran SK, Loving CL. Regional epithelial cell diversity in the small intestine of pigs. Journal of Animal Science. 2023;101:skac318. 10.1093/jas/skac318

31. Nio-Kobayashi J, Takahashi-Iwanaga H, Iwanaga T. Immunohistochemical Localization of Six Galectin Subtypes in the Mouse Digestive Tract. J Histochem Cytochem. 2009;57:41–50. 10.1369/jhc.2008.952317

32. Thompson CA, Wojta K, Pulakanti K, Rao S, Dawson P, Battle MA. GATA4 Is Sufficient to Establish Jejunal Versus Ileal Identity in the Small Intestine. Cellular and Molecular Gastroenterology and Hepatology. 2017;3:422–46. 10.1016/j.jcmgh.2016.12.009

33. Kayisoglu O, Weiss F, Niklas C, Pierotti I, Pompaiah M, Wallaschek N, et al. Location-specific cell identity rather than exposure to GI microbiota defines many innate immune signalling cascades in the gut epithelium. Gut. 2021;70:687–97. 10.1136/gutjnl-2019-319919

34. Goto Y, Uematsu S, Kiyono H. Epithelial glycosylation in gut homeostasis and inflammation. Nat Immunol. 2016;17:1244–51. 10.1038/ni.3587

35. Galeev A, Suwandi A, Cepic A, Basu M, Baines JF, Grassl GA. The role of the blood group-related glycosyltransferases FUT2 and B4GALNT2 in susceptibility to infectious disease. International Journal of Medical Microbiology. 2021;311:151487. 10.1016/j.ijmm.2021.151487

36. Hamdan FH, Farhadipour M, Perez I, Kossick K, Smith H, Edwinson A, et al. Intestinal Stem Cells From Patients With Inflammatory Bowel Disease Retain an Epigenetic Memory of Inflammation. Cellular and Molecular Gastroenterology and Hepatology. 2026;20:101774. 10.1016/j.jcmgh.2026.101774

37. Menche C, Farin HF. Strategies for genetic manipulation of adult stem cell-derived organoids. Exp Mol Med. 2021;53:1483–94. 10.1038/s12276-021-00609-8

38. Wells MF, Nemesh J, Ghosh S, Mitchell JM, Salick MR, Mello CJ, et al. Natural variation in gene expression and viral susceptibility revealed by neural progenitor cell villages. Cell Stem Cell. 2023;30:312–332.e13. 10.1016/j.stem.2023.01.010

39. Adeniyi-Ipadeola GO, Hankins JD, Kambal A, Zeng X-L, Patil K, Poplaski V, et al. Infant and adult human intestinal enteroids are morphologically and functionally distinct. Coyne CB, editor. mBio. 2024;15:e01316–24. 10.1128/mbio.01316-24

40. UE 3P. Pig Physiology and Phenotyping Experimental Facility [Internet]. INRAE; 2018 [cited 2026 July 31]. 10.15454/1.5573932732039927E12

