## Supplementary Figure for "Adult porcine intestinal organoids as models for regional epithelial identity and individual regulatory variation"

### Slide 1
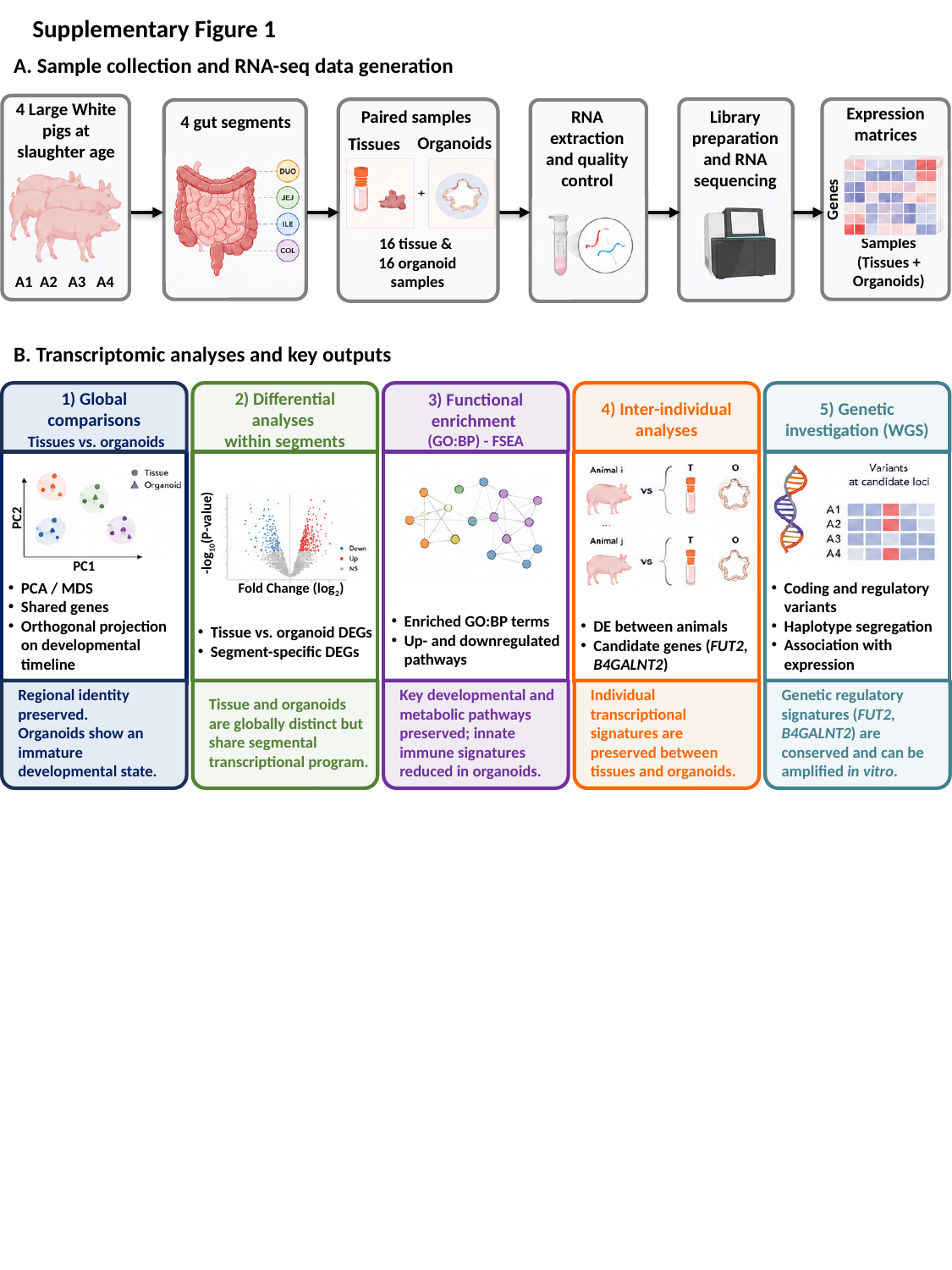

Supplementary Figure 1
A. Sample collection and RNA-seq data generation
4 Large White pigs at slaughter age
Expression matrices
Paired samples
Library preparation and RNA sequencing
RNA extraction and quality control
4 gut segments
Organoids
Tissues
Genes
Samples
(Tissues + Organoids)
16 tissue &
16 organoid samples
A1 A2 A3 A4
B. Transcriptomic analyses and key outputs
1) Global comparisons
 Tissues vs. organoids
2) Differential analyses
within segments
3) Functional enrichment
(GO:BP) - FSEA
4) Inter-individual analyses
5) Genetic investigation (WGS)
PC2
-log_10_(P-value)
PC1
PCA / MDS
Shared genes
Orthogonal projection on developmental timeline
Coding and regulatory variants
Haplotype segregation
Association with expression
Fold Change (log_2_)
Enriched GO:BP terms
Up- and downregulated pathways
DE between animals
Candidate genes (FUT2, B4GALNT2)
Tissue vs. organoid DEGs
Segment-specific DEGs
Regional identity preserved.
Organoids show an immature developmental state.
Tissue and organoids are globally distinct but share segmental transcriptional program.
Key developmental and metabolic pathways preserved; innate immune signatures reduced in organoids.
Individual transcriptional signatures are preserved between tissues and organoids.
Genetic regulatory signatures (FUT2, B4GALNT2) are conserved and can be amplified in vitro.

### Slide 2
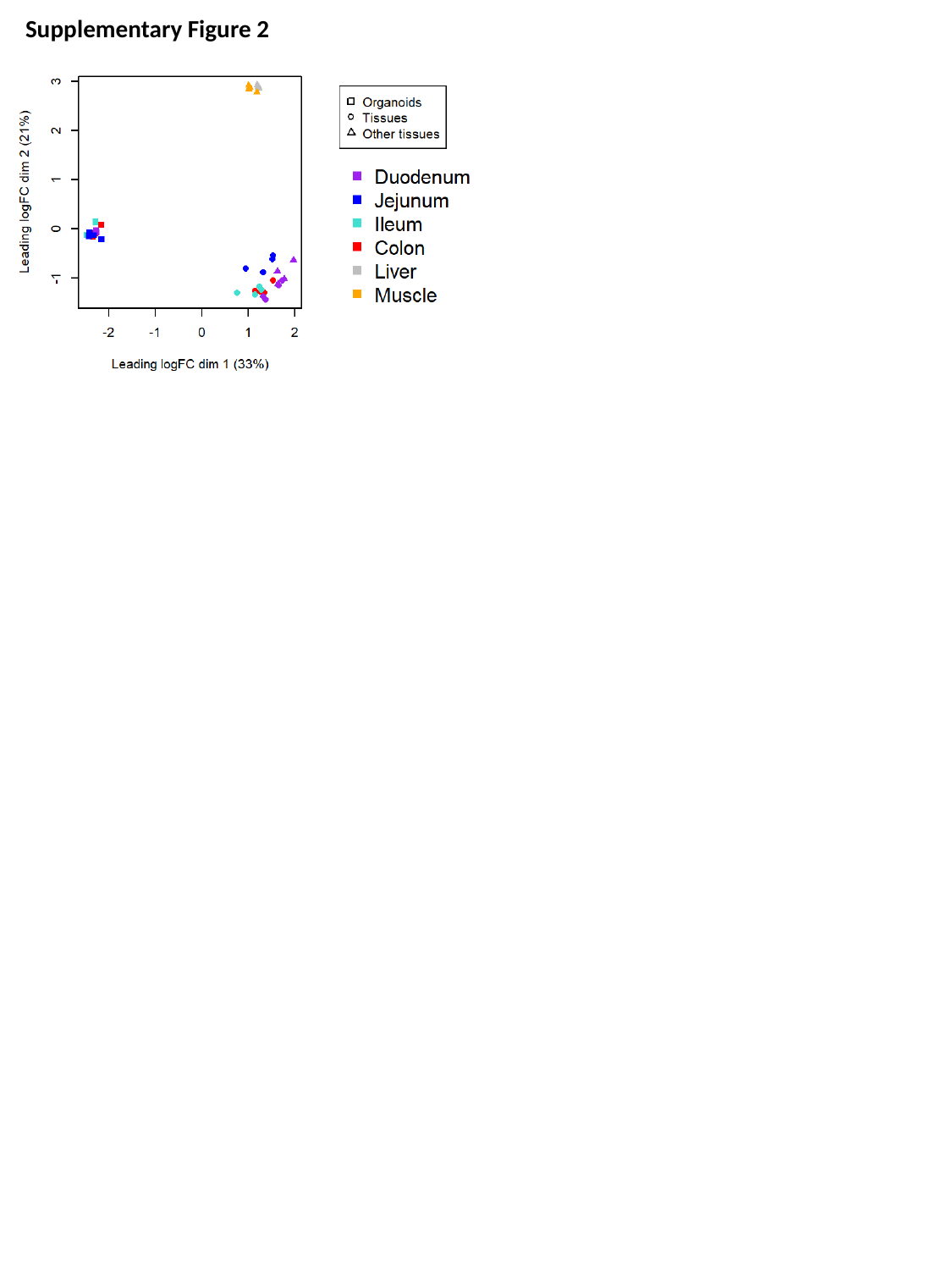

Supplementary Figure 2

### Slide 3
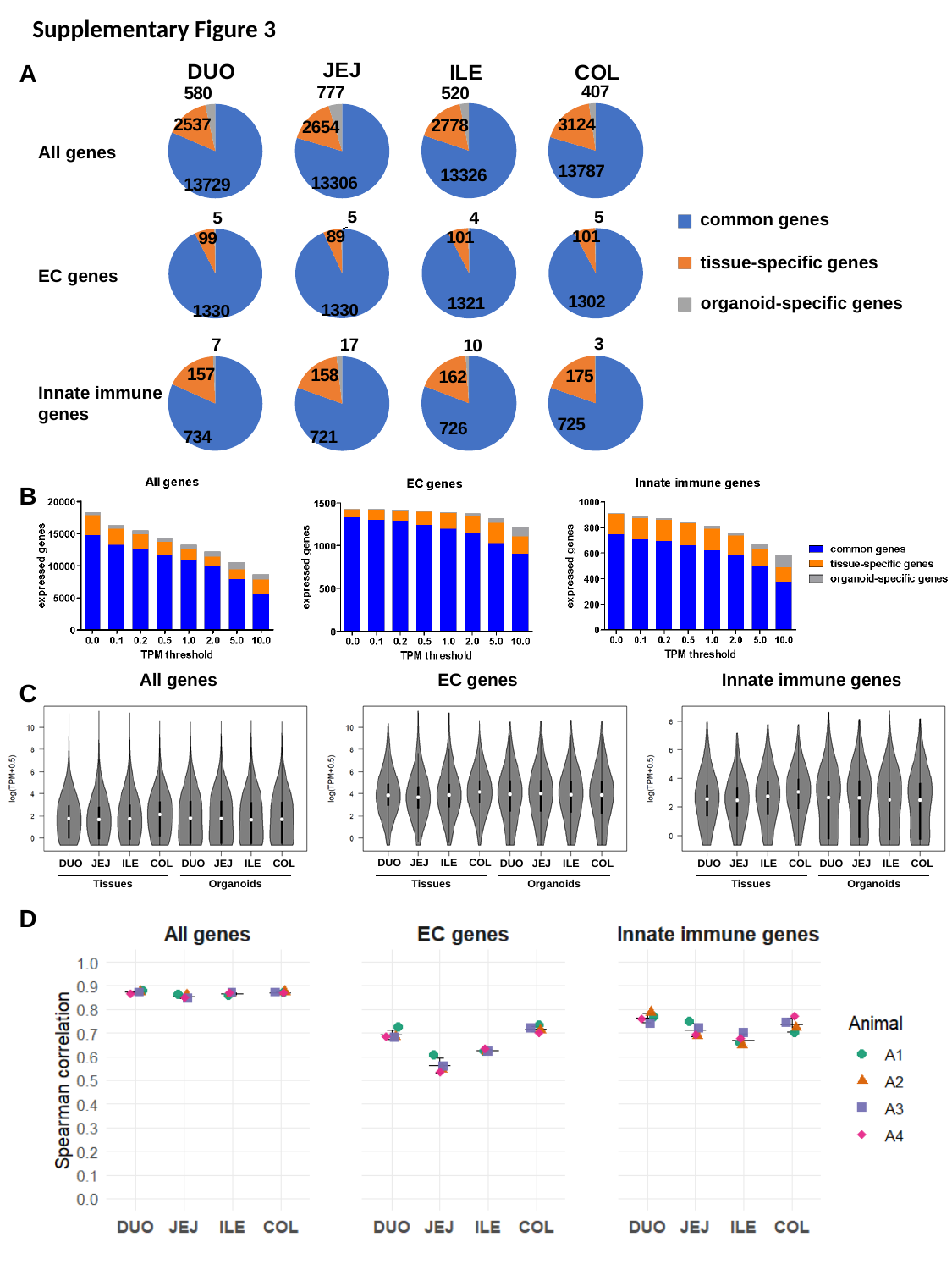

Supplementary Figure 3
A
#### Chart:
| Category | ILE |
|---|---|
| common genes | 13326.0 |
| tissue-specific genes | 2778.0 |
| organoid-specific genes | 520.0 |
#### Chart:
| Category | COL |
|---|---|
| common genes | 13787.0 |
| tissue-specific genes | 3124.0 |
| organoid-specific genes | 407.0 |
#### Chart:
| Category | DUO |
|---|---|
| common genes | 13729.0 |
| tissue-specific genes | 2537.0 |
| organoid-specific genes | 580.0 |
#### Chart:
| Category | JEJ |
|---|---|
| common genes | 13306.0 |
| tissue-specific genes | 2654.0 |
| organoid-specific genes | 777.0 |All genes
#### Chart
| Category | ILE |
|---|---|
| shared | 1321.0 |
| specific T | 101.0 |
| specific O | 4.0 |
#### Chart
| Category | COL |
|---|---|
| shared | 1302.0 |
| specific T | 101.0 |
| specific O | 5.0 |
#### Chart
| Category | DUO |
|---|---|
| shared | 1330.0 |
| specific T | 99.0 |
| specific O | 5.0 |
#### Chart
| Category | JEJ |
|---|---|
| shared | 1330.0 |
| specific T | 89.0 |
| specific O | 5.0 |common genes
tissue-specific genes
EC genes
organoid-specific genes
#### Chart
| Category | ILE |
|---|---|
| shared | 726.0 |
| specific T | 162.0 |
| specific O | 10.0 |
#### Chart
| Category | COL |
|---|---|
| shared | 725.0 |
| specific T | 175.0 |
| specific O | 3.0 |
#### Chart
| Category | DUO |
|---|---|
| shared | 734.0 |
| specific T | 157.0 |
| specific O | 7.0 |
#### Chart
| Category | JEJ |
|---|---|
| shared | 721.0 |
| specific T | 158.0 |
| specific O | 17.0 |Innate immune genes
B
All genes
EC genes
Innate immune genes
DUO JEJ ILE COL
DUO JEJ ILE COL
DUO JEJ ILE COL
DUO JEJ ILE COL
DUO JEJ ILE COL
DUO JEJ ILE COL
Tissues
Organoids
Tissues
Organoids
Tissues
Organoids
C
D

### Slide 4
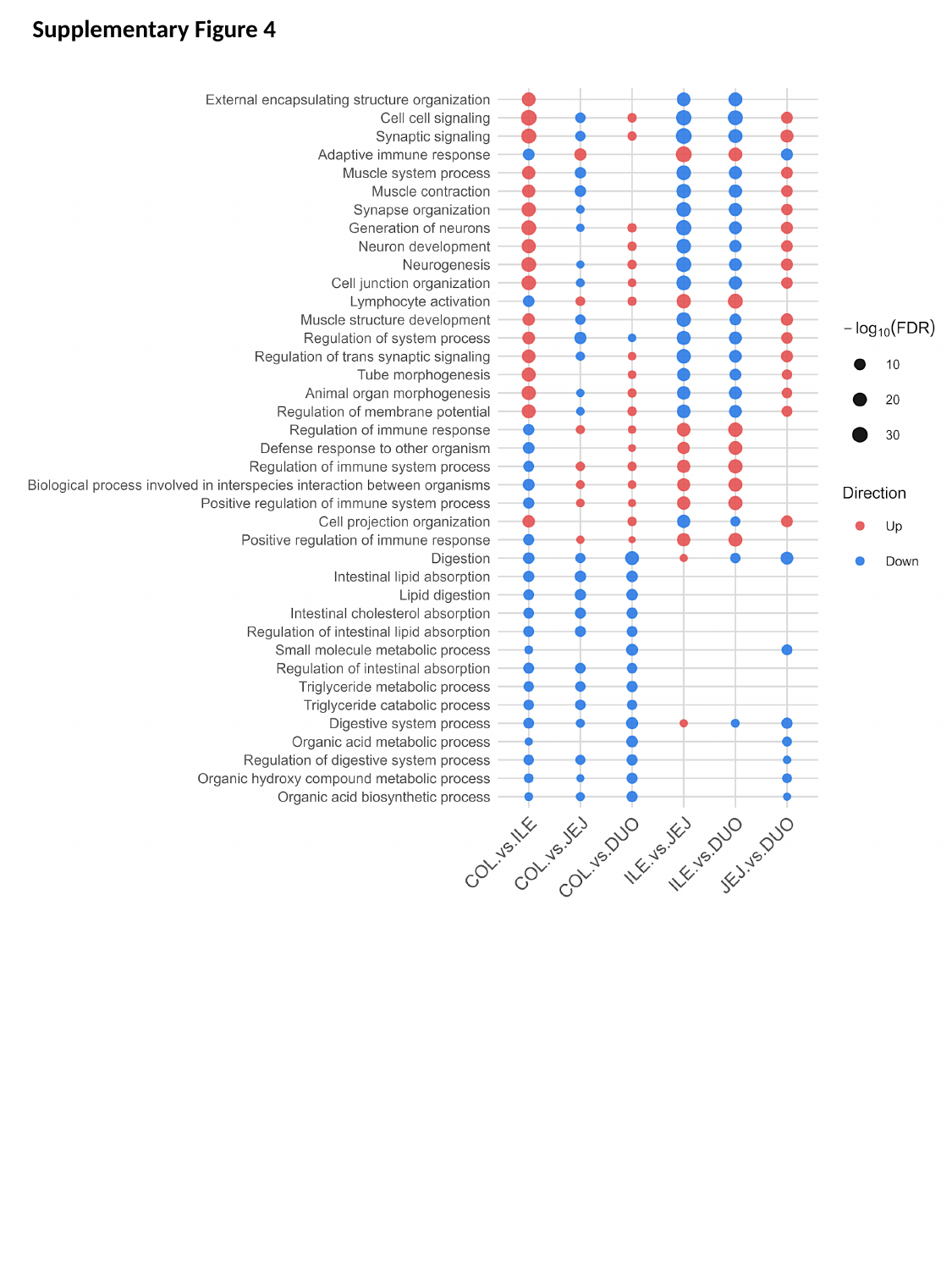

Supplementary Figure 4

### Slide 5
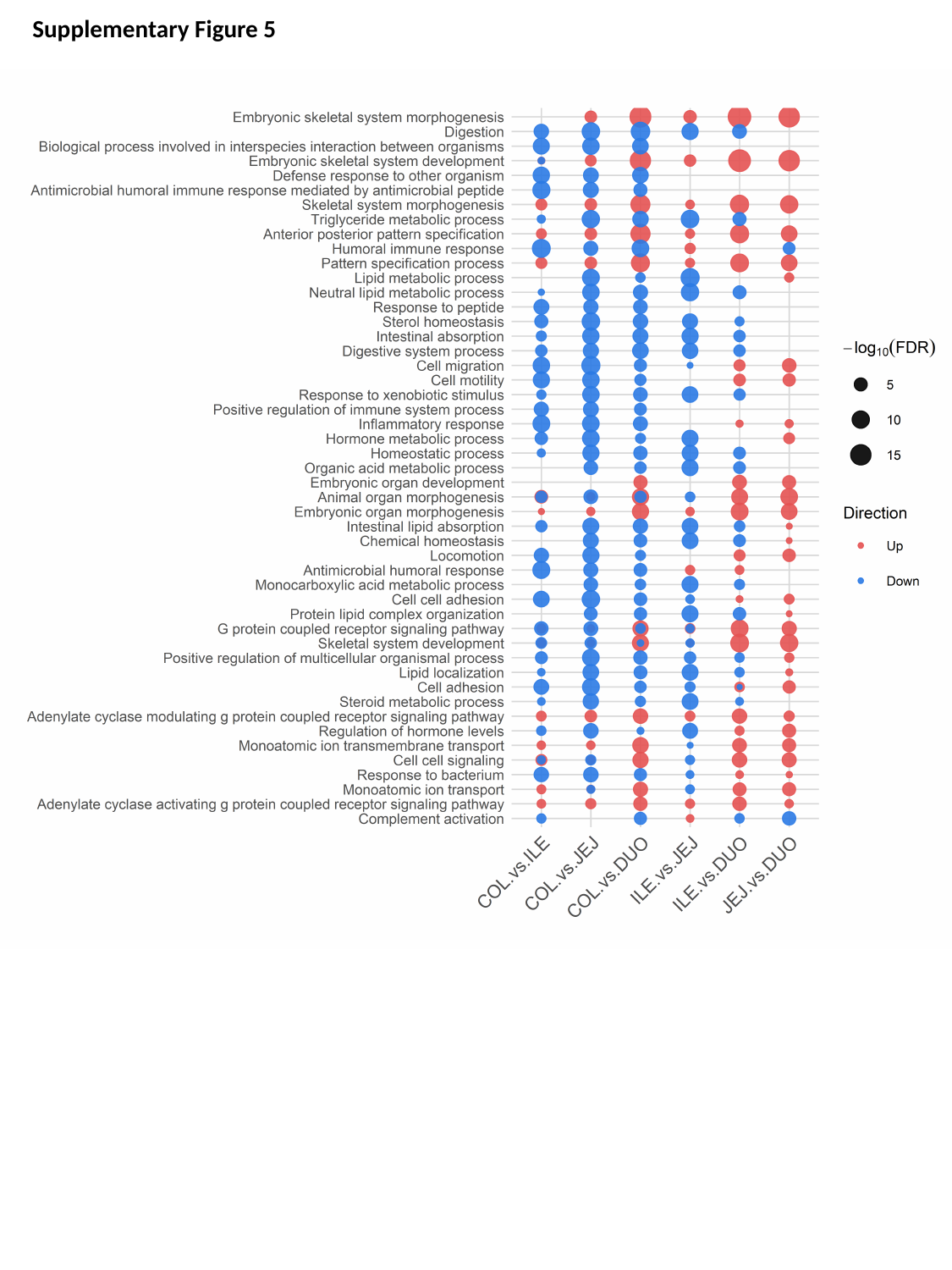

Supplementary Figure 5

### Slide 6
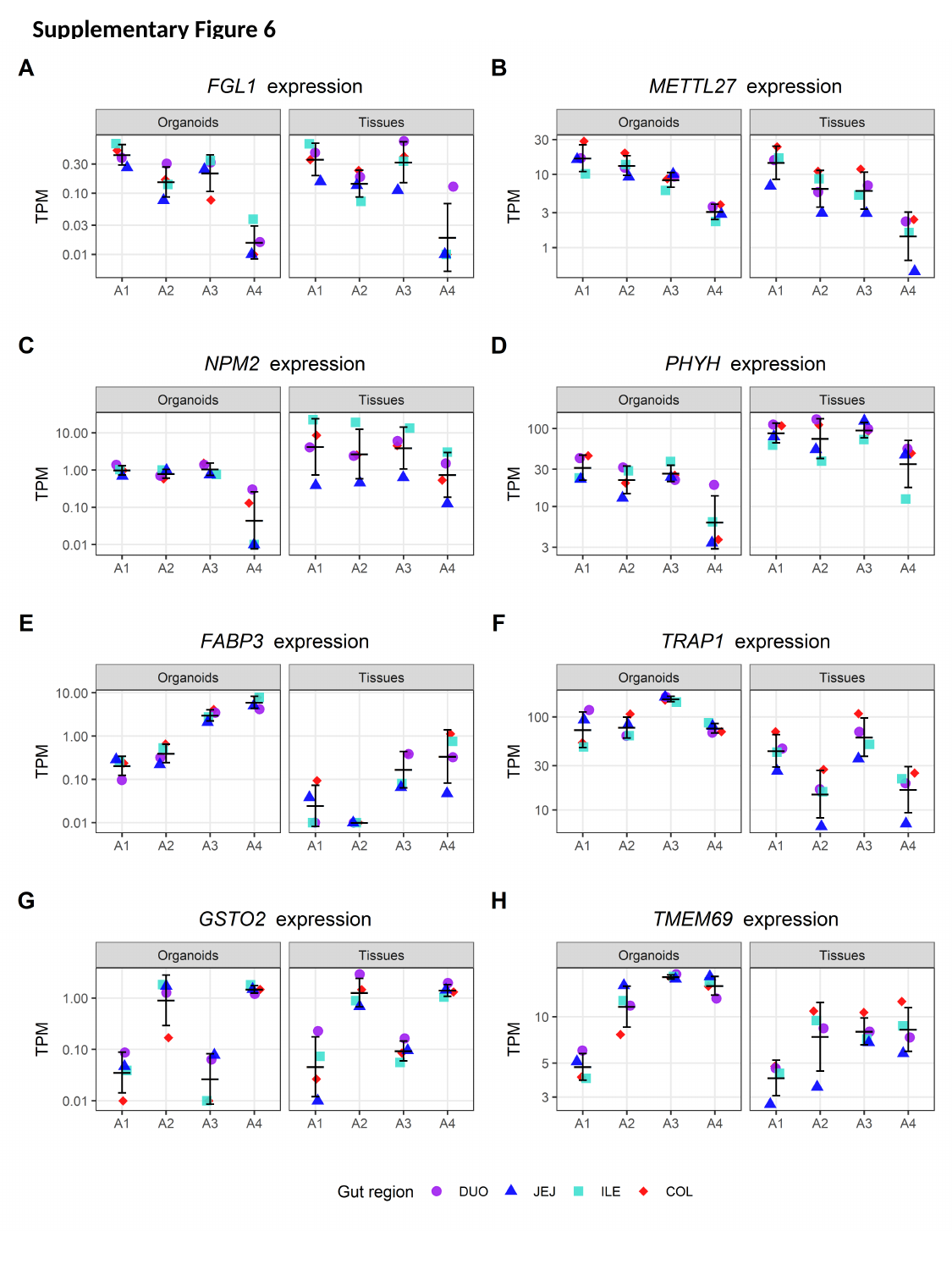

Supplementary Figure 6

### Slide 7
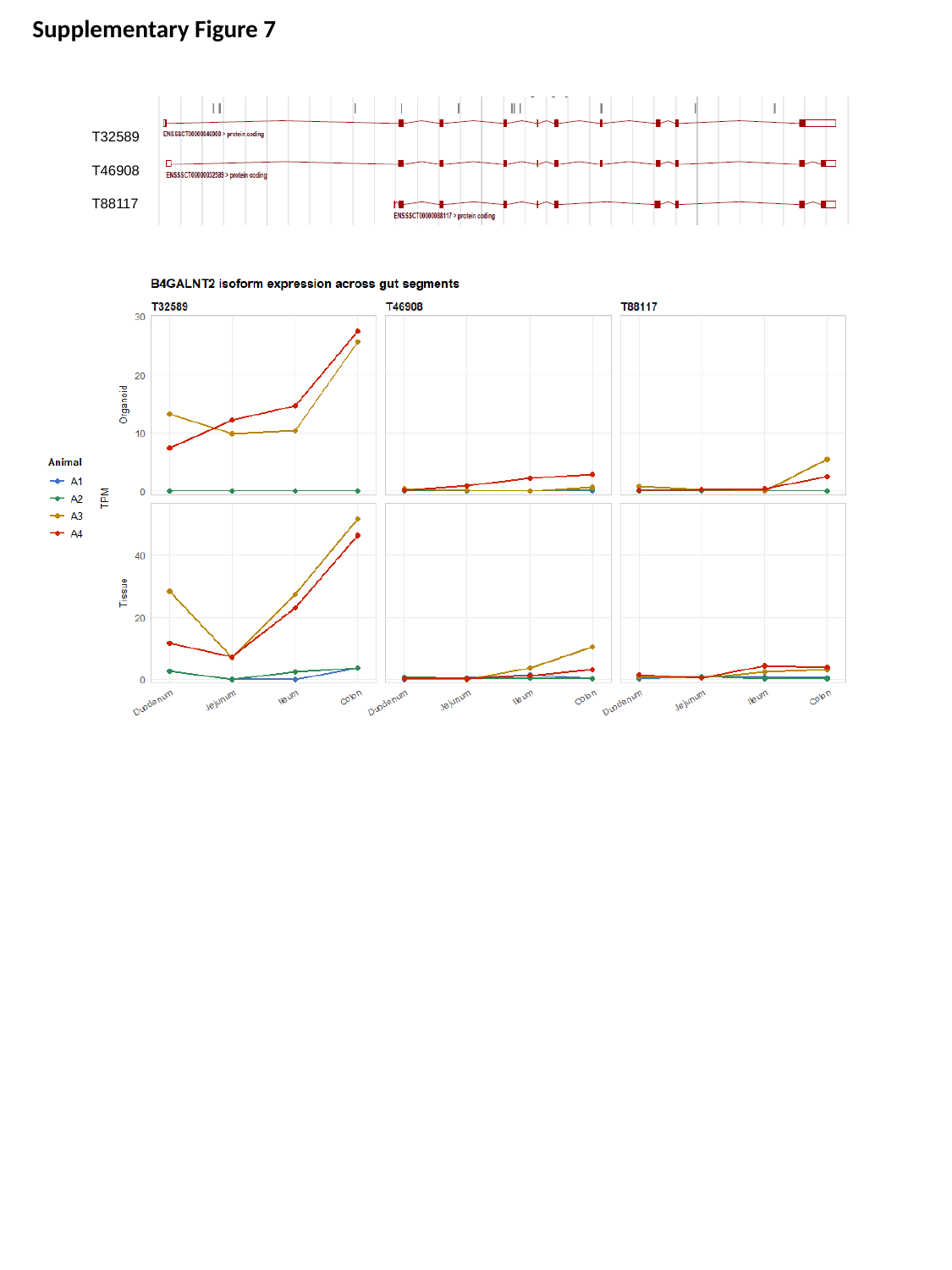

Supplementary Figure 7
T32589
T46908
T88117
